# Graph-Based Pharmacokinetic-Pharmadynamic Modeling for Large Scale Systems: Nanoparticles Case

**DOI:** 10.1101/2022.07.12.499805

**Authors:** Teddy Lazebnik, Hanna Weitman, Gal A. Kaminka

## Abstract

Pharmaceutical nanoparticles (NPs) carrying molecular payloads are used for medical purposes such as diagnosis and medical treatment. They are designed to modify the pharmacokinetics-pharmacodynamics (PKPD) of their associated payloads, to obtain better clinical results. Currently, the research process of discovering the PKPD properties of new candidates for efficient clinical treatment is complicated and time-consuming. *In silico* experiments are known to be powerful tools for studying biological and clinical processes and therefore can significantly improve the process of developing new and optimizing current NPs-based drugs. However, the current PKPD models are limited by the number of parameters they can take into consideration and the ability to solve large-scale *in vivo* settings, thus providing relatively large errors in predicting treatment outcomes. In this study, we present a novel mathematical graph-based model for PKPD of NPs-based drugs. The proposed model is based on a population of NPs performing a directed walk on a graph describing the blood vessels and organs, taking into consideration the interactions between the NPs and their environment. In addition, we define a mechanism to perform different prediction queries on the proposed model to analyze two *in vivo* experiments with eight different NPs, done on mice, obtaining a fitting of 0.84 ± 0.01 and 0.66 ± 0.01 (mean ± standard deviation), respectively, comparing the *in vivo* values and the *in silico* results.

## 1 Introduction

The investigation and discovery of new medical nanoparticles (NPs) is an active field of study [33]. In 2016 alone, more than 50 NP-based medical treatments have been approved by the food and drug administration (FDA) [16]. Improving and accelerating the investigation of medical NP raises many challenges [19].

Mathematical models and computer simulations are effective tools to investigate clinical treatments [2]. Specifically, Pharmacokinetics-Pharmacodynamics (PKPD) models have been investigated for multiple medical treatments and environments [14, 52, 15, 61] and shown to well represent targeted drug delivery [31]. *In silico* experiments accelerate the development and testing processes of new medicines and contribute to our understanding of the dynamics between the body and the treatment [26, 21, 37]. However, since the mathematical approaches currently used as the framework of the PKPD models are mainly based on cells populations and chemical interactions, the exciting models are mathematically complex and require significant computation. Therefore, they simplify some parameters of the biological systems by making assumptions that may be invalid [51]. For example, the cardiovascular system is often treated as a single organ and modeled with homogeneous behavior all over its geometry [18, 42].

We propose a novel approach to modeling based on a graph representation of the blood vessels and organs. A population of nanoparticles (NPs) performing a directed walk on this graph, interacting with itself and with the environment (i.e., the blood vessels and organs). This approach allows accurate simulations even for large-scale *in vivo* systems. Based on the proposed model and on *in vivo* data, we developed a simulator implementing the proposed model. We reproduced the *in vivo* values of two NP’s families [12, 40] using the simulator. They have six and two types of NPs, respectively. We simulated each of the NPs, comparing the *in silico* results to the *in vivo* values presented by Ben-Akiva et al. [12] and Lee et al. [40], respectively. We successfully demonstrated that the proposed model reproduces the *in vivo* values of two different studies, linear regression of the *in silico* experiment results in respect to the *in vivo* values were 0.842 (*R*^2^ = 0.933) and 0.659 (*R*^2^ = 0.946), respectively.

This paper is organized as follows: In section 2, we present an overview of PKPD modeling in general and both formula-based and ordinary differential equation (ODE) based PKPD models, in particular. Afterward, biological graph-based mathematical models and the transformation between graph-based and ODE-based representations of biological systems are described. In Section 3, we proposed a new mathematical model with an analytical formalization of computing predictions using the proposed model, inspired by NP-based drug dynamics. In Section 4, we present how to numerically and analytically answer any prediction query. In Section 5, a simulation based on a mouse’s anatomy with two NPs families is presented. A comparison between the simulation’s (*in silico*) results and the *in vivo* values is provided. Finally, in Section 6, we discuss the main advantages and limitations of the model compared to other PKPD models, and in Section 7 conclude the contribution of this work and propose future work.

## 2 Related Work

Medical NP is a promising growing approach that is used for a wide range of medical purposes like diagnosis, monitoring, and treatment; to name a few. The process of investigating and discovering new medical NPs is complex, time-consuming, and requires researchers from various disciplines like chemistry, biology, and physics [41, 23]. NPs cover a wide range of materials whose dimensions are in the nanometric scale between 1 to 100 nm. They have novel properties and functions due to their nanometric size [34]. A major advantage of medical NPs is their ability to efficiently carry and deliver drugs to specifically targeted sites and release them in a controlled manner as drugs are loaded on NPs and injected into the bloodstream.

Medical NPs are designed to modify the PKPD of their associated drugs to overcome the physiological barriers to efficient drug delivery [11, 24, 45]. To develop and test new drug-carrying NPs for targeted drug delivery, and to better understand their dynamics in the body, one can take advantage of *in silico* experiments which are based on mathematical models of the biological and clinical processes [2, 26, 21, 37], as conducted in this research. For this, we must develop a highly accurate representation of the dynamics of medical NPs *in vivo*.

Clinical treatment involves the administration of NPs by intravenous injection of single- or multiple-doses of NPs into the bloodstream. It is known that NPs concentration decreases over time and that NPs’ half-life depends on their location [5]. The NPs are carried in the bloodstream and accumulate in different organs and tissues in the body. The cardiovascular system can be described as a graph where the connections between either multiple blood vessels and a blood vessel or an organ define both the blood vessel itself and to what it is connected.

NPs’ life span in an organ is affected by its role and its tissue microenvironment which is different for each organ [22, 8]. For example, some of the NPs are filtered from the blood circulation system by the liver, spleen, and adrenal. They can biodegraded, release their payload, react with each other, aggregate, etc [47, 4]. Based on that, several works [53, 28, 29] proposed that each NP can be described using a finite number of states with some physical and chemical properties. For example, a DNA origami NP that contains a payload may have three states: close with payload, close without payload, and open [3].

A review of PKPD models is presented, including Formula-based and ODE-based models with the pros and cons of each one of them as methods for large-scale *in silico* experiments. Afterward, several graph-based mathematical models for biological systems are presented, showing studies where using this representation allowed us to investigate large-scale biological systems.

### 2.1 Pharmacokinetics-Pharmacodynamics models

Pharmacokinetics (PK) and Pharmacodynamics (PD) models are a group of models that describe drug interactions with living organisms and vice versa, respectively [55]. In particular, PK models are designed to describe the absorption, distribution, metabolism, and elimination process of drugs while PD models deal with the effect of the drug on the body and the rate of change in the drug concentration in the bloodstream. Integration of these two models is referred to as PKPD models.

PKPD models have been investigated for multiple NP and organs tissues microenvironments [**new·nano·pkpd**, 31, 58] and shown to well represent drug delivery [31]. Nonetheless, these models are focused on parts of the biological system or simplify some parameters of the biological systems by making assumptions that may be shown to be invalid [51] due to the mathematical complexity associated with representing this kind of dynamics. For example, the cardiovascular system is modeled as a single organ and therefore acts homogeneously all over its dynamics. Some assumptions are commonly used in the PKPD models to handle the mathematical and computational complexity associated with modeling biological systems.

In order to reduce the number of assumptions and better represent a biological system, a model should take into consideration a large number of processes while keeping the computation necessary for either numerically or analytically obtaining the model’s state feasible. The common mathematical representation for PKPD models can be divided into two main groups: Formula-based and ODE-based. Each group has advantages and disadvantages regarding the ability to accurately represent biological systems and the computation required to compute.

#### 2.1.1 Formula-based PKPD models

Formulas-based PKPD models are functions getting one or more biological parameter and returning another biological quanta [36, 60, 32]. Formally, a formula-based PKPD model can be desired by a function *F* : ℝ^*ν*^ → ℝ, where *ν* ≥ 1. For instance, Chandasana et al. [20] suggested that a NP-based drug dissolve in a body is defined by the formula

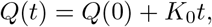

where *Q*(*t*) is the amount of drug dissolved at time *t, Q*(0) is the initial amount of drug in the solution (e.g., *t* = 0), and *K*_0_ is the zero order release constant. In this case, one set the desired time point of the treatment and gets the drug dissolve in the body. The parameters of the formulas-based PKPD models usually fitted to either *in vivo* or *in vitro* experimental results [36, 60].

The advantage of these models is their simplicity which allows both analytical analysis of the system’s dynamics and fast calculations. The calculations required for this type of model grow slowly as a function of the model’s size while providing results with similar accuracy to more advanced methods. The growth of the computation required to solve the model as a function of the number of data points in the biological model is significantly smaller in the group of formula-based PKPD models, compared to our model. However, these models can answer only one prediction query which they were specifically designed for. To answer another query, a new model must be developed from scratch which again requires mathematical and biological expertise. Our model shows a significant advantage by allowing infinite prediction queries.

#### 2.1.2 ODE-based PKPD models

Differential-equations-based PKPD models are represented by a system of ordinary or partial differential equations (ODEs/PDEs) which represent the changes in the dynamic of the system over time. The system is solved numerically or analytically to arrive at predictions.

Wei-Yu et al. [62] proposed a PK model to evaluate the accumulation of various sizes of *ZnO* NPs in tissues of mice over time. It is easily computed by their asymptotic solution. Wei-Yu et al. showed that their model after seven simulated days barely fit the experimental data for one NP while better approximating the experimental data for another NP [62]. The authors claim the error originated in a process the model does not take into consideration as it takes place in an organ that is not included. To overcome similar errors, a larger model enables us to take more organs and processes into consideration is required. It is possible to extend the current model but this requires revising all of the equations. In contrast, a graph-based approach allows for local changes without influencing the entire model.

Kristensen et al. [39] proposed a method that is based on stochastic differential equation (SDE) and in vivo data. The method is able to produce a system of non-linear SDEs to describe data using eight-step fitting processes developed by the authors which lie on a direct search for the equations structure and variables values throughout a trial and error process. The authors used their model on a simple absorption kinetics case, showing an accurate approximation of *in vivo* data. However, this method is able to represent a single biological process, and assumes a maximum of each SDE in the model is either first or second order. Therefore, the model is limited to representing a single biological process at a time and cannot handle the complexity of representing whole-body dynamics.

Friebiesm et al. [9] outline a PKPD model to quantify physiologic resistance of tumor tissue growing to drugs in a three-dimensional, *in vitro* case using a system of three second-order, non-linear PDEs that describe the spatio-temporal grows of the tumor and taking into consideration the interaction with the injected drug. The authors analyzed the *in vitro* case rather than the *in vivo* case in order to ignore the role of the metabolism in the dynamics, allowing the authors to calculate tumor drug response as a predictable process dependent on biophysical laws. Indeed, Friebiesm et al. [10] outline the additional complexity associated with the heterogeneous infiltrative morphologies which depend on the magnitude of cell adhesion forces. In comparison, one can use a graph-based approach to divide the geometrical configuration into small, homogeneous regions, solving for a similar dynamics of the *in vitro* case and adding interaction between regions, obtaining a feasibly numerically solvable model.

Namazi et al. [46] proposed a phase lagging model for drug diffusion in solid tumors describes the relationship between the drug concentration in the solid tumor and its spatial-temporal effect. The authors developed a non-linear, first-order PDE model, taking into consideration one spatial dimension with diffusion and drug penetration dynamics. The authors stated that the proposed model’s accuracy can be significantly improved by integrating additional biological factors (for instance, cancer spread) however given the current representation of the model such introduction would require re-developing the entire model.

These models take into consideration several to a few dozens of processes and in-vivo locations (either organs, tissues, or cancer tumors) because large-scale ODE/PDE systems are difficult to analyze. Furthermore, *in vivo* analysis is a result of a nonlinear system that is considered complicated to explicitly analyze [57, 25].

Overall, numerical-based (either formula- or ODE-based) models well perform on relatively small-size biological models and for a relatively short period. For example, one blood vessel (5 seconds) [43] and 84 blood vessels (0.14 seconds) [7]. In these scales, numerical-based models are able to provide accurate results [43, 7]. Nevertheless, these models are inherently introducing approximation error which increases over simulation time or in correlation to the model’s size. As a result, currently, numerical models do not scale well and therefore can hardly handle the complexity of large-scale biological systems.

### 2.2 Biological graph-based models

Graph-based models for biological systems are commonly used as an effective modeling approach for complex and large scale systems [59, 56, 38, 50, 1]. Shih-Yi reviewed the usage of graph-based modeling for cell biology network systems, such as protein-protein interactions and metabolic network [56]. The authors show that graph-based models can exploit global and local characteristics of biological networks relevant to cell biology in multiple cases but conclude that the transitional graph-based model is usually not sophisticated enough for effective treatment strategies for diseases such as cancer [56].

Tashakor and Suppi [59] investigated an agent-based tumor model where each node in the graph is a cancer cell with a finite state and multiple biological features and the edges are the biological connections between the cells. The graph changes its topology over time as a result of several biological stochastic processes. The authors use a graph to represent interactions between cells while we use graphs to represent the physical connection between (blood vessels and organs. Namely, our model represents physical locations in which agents (NPs) can interact and as such inherently defines a time-depended interaction graph like the one proposed by Tashakor and Suppi [59]. Hence, we proposed an approach that generates a time-depended interaction graph rather than directly defining one.

Similar to our approach, Reichold et al. [35] investigated the vascular graph with blood flow where nodes represent locations in which vessels bifurcate or end, and edges designate the vessels themselves. The authors’ blood flow model was defined as a function of blood pressure differences between the nodes. Blood flow velocity depends on the pressure differences and the conductance of the vessels and was modeled by using linear first-order ODE. The model contains information on the graph in a 3-dimensional space. In comparison, we model the blood vessels as nodes and the connections between them resulting from bifurcations as edges. This allows getting a representation of a blood vessel or part of it by replacing one node with the corresponding line graph of this node. In addition, our approach does not take into consideration the 3-dimensional geometrical configurations of the nodes which can be used in models considering thermodynamics.

In particular, Poelma [48] investigated the use of a general cardiovascular graph analyzing the pressure and flow rate distribution from either imaging data or prescribed (e.g. a hexagonal lattice). Specifically, Ganesan, He, and Xu [30] investigated the coronary blood flow of a living pig [30]. Popel and Gross [49] investigated the transport of oxygen in arteriolar networks in a hamster. All these model used the graph of the vascular system as the spatial part of the model, computing different types of biological interactions in them. In a similar manner, our model assumes flow dynamics with homogeneous pressure and linear flow of NPs inside the cardiovascular system and several organs.

## 3 A Graph-based PKPD Model

The proposed model is constructed from three main components: a population of NPs containing a payload drug (or operating themselves as a drug), a graph representing abstract physical locations in the body, divided into blood vessels and organs, in which the NPs are located at and moving between due to the blood’s natural flow, and the dynamics that include the NPs interact with other NPs and with the environment at each point in time.

A schematic view of the model’s dynamics is shown in Fig. 1 where there are three locations (blood vessel, organ, blood vessel) with two NPs in the first two locations. During a single step in time, the NPs are flowing between the locations due to the blood’s flow inside the blood vessel and the organ. In addition, *NP*_1_ and *NP*_2_ interacted, causing *NP*_2_ to release its payload. Moreover, *NP*_3_ interacted with the organ, causing it to be removed from the body (for instance, the liver eliminates NPs).

**Figure 1:**
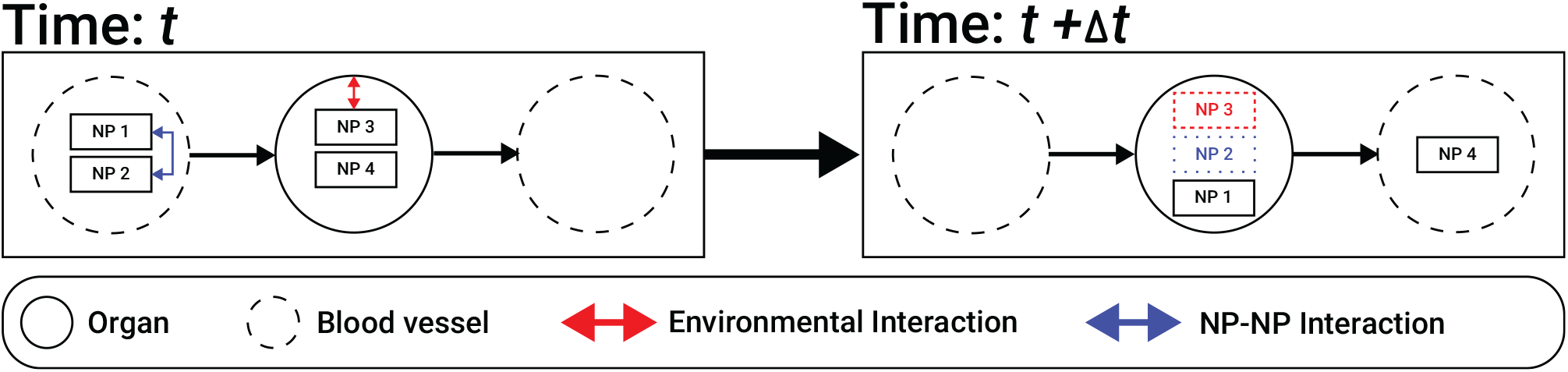
A schematic view of the model’s dynamics.

### 3.1 Model definition

Inspired by the dynamics of the biological systems, we propose a mathematical model allowing analysis and prediction of the behavior of NPs within a biological system. The model is defined by a tuple *M* := (P, *G, I*) where P is a set (the population) of NPs, *G* is a flow graph of the model representing the network of locations where NPs can be found in the body, and *I* is the interaction protocol between the NPs. The components of the tuple are described below in detail.

#### 3.1.1 The population of nanoparticles ℙ

We consider the NPs as agents with one or more states. Mathematically, let ℙ be a non-empty set of NPs such that each *np* ∈ ℙ is defined by a tuple *np* := (*s*(*t*), *A, µ, type*) where *s*(*t*) is the state of the NP at a given time *t, A* := [1, … *a*], *a* ∈ ℕ ^+^ is a finite space of possible states the NP can take such that *s* ∈ *A, µ* ∈ ∅ ∪ 𝔻 is the payload drug the NP holds where 𝔻 is the set of all possible drugs, and *type* is a category of a state machines that the NP related to.

The *µ* at time *t* = 0 can be either ∅ or *D* ∈ 𝔻. The value of *µ* can transform to ∅ from any other value if *s*(*t*) ∈ *Z* ⊂ *A* and *s*(*t* − 1) ∈ *A/Z* ⊂ *A* where *Z* is the set of states that release the drug of the NP. A drug that released from the NP is effective the immediate location the NP is at at time *t*.

The *type* property allows to separate the NPs’ population into sub populations which later be useful dealing with each type of NP’s interaction according to its type and state. 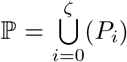 such that each group {*P*_*i*_} for *i*∈ [1, … ζ] is the subset of NPs from the same type.

In addition, we assume that either the NPs’ population ℙ decreases over time as a result of the biodistribution and clearance of the NPs from the bloodstream 𝔽 over time or that each NP has a pre-given life span. In both cases, the distribution and the NP’s life span can be changed due to the interactions of NPs. Formally, *t*_*i*_ > *t*_*j*_ ∈ ℝ^+^ : |ℙ (*t*_*i*_)| < |ℙ (*t*_*j*_)| and that for each NP, the state space (*A*) contains a “inactive” state *ξ* ∈ *A* such that NP can reach due to interactions with other NPs or spontaneously as a function of time.

#### 3.1.2 The flow graph *G*

Let *G* be a graph defined by the tuple *G* := (*V, E*), where *E* ⊂ *V* × *V* and *V* := *O* ∪ *B* such that *O* ∩ *B* = ∅. *O* is the set of organs in the body, *B* is the set of blood vessels, and *E* is the edges between organs, blood vessels, and themselves.

Each organ node *o* ∈ *O* defined by a tuple *o*(*t*) := (*P* (*t*), *ts*, Ψ, *d*) for a given point in time *t*, where *P* (*t*) ⊆ ℙ is the population of NPs inside the organ in a given time, *ts* > 0 ∈ ℕ is the number of clocks tics a NP inside the organ (*np* ∈ *P* (*t*)) is staying inside the organ before it moves to another node, Ψ is a stochastic function Ψ : *P* (*t*) → *P* (*t*) changing the states of each NP *np* ∈ *P* (*t*) with a probability *α* ∈ [0, 1] × …_*ζ*_ × [0, 1] or make the NP stay inside this organ node (*o* ∈ *O*) from this step forward depending on the NP’s type, and *d* is a function *d* : 𝔻^*n*^ → ℝ^*n*^ where 𝔻 is the set of all drugs in the system mapped to the amount of each drug in the node *d* : *P* (*t*) → *P* (*t*).

*B* are the blood vessels in the body, in which we assume NPs do not have any interaction and are used for the flow of the NPs’ population. Therefore, each blood vessel node *b* ∈ *B* is a private case of an organ node *o* such that *ts* = 1, Ψ = *Id*, and *d* = ∅. Of note, *G* is a simple graph which means it does not contain loops (i.e., *e* = (*v*_*i*_, *v*_*i*_)) or nodes with more than one edge between them.

Finally, We define *E* as the set of edges between the different nodes (*V*). Each *e* ∈ *E* defined by the tuple *e* := (*v*_1_, *v*_2_, *w*), where *v*_1_, *v*_2_ ∈ *V* are the source and target node, and *w* ≥ 0 ∈ ℕ is the absolute weight represents how many NPs can move along this edge in a single clock tick. To represent close cardiovascular systems with connected organs, we assume *G* is a connected graph. In addition, we assume that *e* = (*v*_*i*_, *v*_*j*_) ∈ *E* ↔ *v*_*i*_, *v*_*j*_ ∈ *B* ∨ *v*_*i*_, *v*_*j*_ ∈ *O*.

#### 3.1.3 Nanoparticles interaction protocol *I*

The NPs’ interact with each other and react to the organ’s tissue on the individual level and at the population level. The population-level interaction is modeled by an interaction protocol that is based on the NPs population and its distribution in the flow graph (*G*). Mathematically, let *I* be an interaction protocol (IP) computational method described by a function *I* : ℙ × *G* → ℙ. In the scope of this model, we define an IP to perform on any set of *k* > 0 NPs, in each interaction. An interaction can change the NP’s state or keep the NP on the same node (*o* ∈ *O*) from this step forward. Furthermore, the IP is stochastic in the manner that an interaction may be performed in *β* ∈ [0, 1] × …_*k*_ × [0, 1] probability for any set of *k* NPs.

Specifically, *I* defines two sub-interaction protocols influencing the NPs’ population size: *L*_*O*_, *L*_*B*_. *L*_*O*_ is a function *L*_*O*_ : {ℙ, *O*} → ℙ which defines the decrease in the NP population currently located in organ nodes for a single step in time. In particular, *L*_*O*_ handles the decrease of the population of NPs in the organs according to the population’s size and NPs’ types in each organ node *o* ∈ *O* separately. Similarly, *L*_*B*_ is aa function *L*_*B*_ : {ℙ, *B*} → ℙ which defines the decrease n the NP population currently located in blood vessel nodes for a single step in time.

The model has a synchronized clock. In the beginning (*t*_0_), the population of NPs ((ℙ) is allocated to one blood vessel (*b*_0_). Then, in each clock tick (marked by *t*_*i*_ for the *i*_*th*_ tic) the following happens: first, for each node *v* ∈ *V* a portion *φ*(*v* → *v*_*i*_) of the NPs population in this node *p* ∈ *v* is moving to a node *v*_*i*_ ∈ *N* (*v*), such that

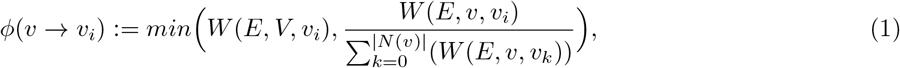

where *W* (*E, v*_1_, *v*_2_) is a function getting a set of edges *E* and two nodes *v*_1_, *v*_2_ and return the weight (*w*) of the edge *e* such that *v*_1_, *v*_2_ are the source and target nodes of the edge, respectively. If no such edge exists, the function returns 0. Namely, the number of NPs that can flow from node *v* to a neighbor node *v*_*i*_ is defined by Eq. (1). The capability of NPs of the specific neighbor node *v*_*i*_ from all the neighbor nodes of node *v* (*N* (*v*)) as a linear approximation of the blood vessels pressure. As an edge case, if the amount of NPs that need to move to the target node *v*_*i*_ is larger than the volume of the target node can store, the maximal number of NPs that can be stored in the target node is moved into it while the rest kept in the same node *v*.

Afterward, in each organ (*o* ∈ *O*) the Ψ interaction protocol is performed. Finally, for each population of NPs (*p* ∈ *v*) the interaction protocol between the NPs (*I*) is performed.

### 3.2 Queries: using the model for analysis

Given a model, one can answer prediction (forward) queries about the modeled system. For example, we may be interested in how long it would take for some amount of gold NPs of a specific type to reach a cancer site. To do this, we use a query on the model with the settings of the experiment and the population of NPs. Based on the results, we can either change something in the NPs to improve the results or reevaluate the results *in vivo* if the *in silico* results are promising.

Each state *S* in time *t* of the model (*S*(*t*)) described above is represented as follows:

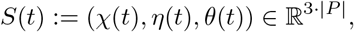

where *χ*(*t*) is the amount of NPs in each node *v* ∈ *V* of the flow graph at time *t, η*(*t*) is the states of the NPs in the population ℙ at time *t*, and *θ*(*t*) is the life span of the NPs in the population ℙ at time *t*.

A *forward query* is a function *FQ* such that *FQ* : (*M, S*(*t*_0_), *SC*) → 𝔽 where *M* is the model, *S*(*t*_0_) is the state of the model at time *t*_0_ (e.g. after first injection of NPs), *SC* is a stop condition function *SC* : *S*(*t*) → {0, 1}, and 𝔽 is the space of all functions *F* that satisfies *F* (*M, S*(*t*_0_)) ⊆ *S* (*t*).

Following the sample example, one can define an FQ as follows. First, the model is defined by previous known bio-clinical dynamics between the gold NPs and the body. The flow graph *G* is given as well and can change between species or even between individuals, given the required data. Then, *S*(*t*_o_) will be the injection location of the gold NPs with their biochemical properties such as the population’s half-life. As such, it also defines the NPs population ℙ. Afterward, one can define the stop condition (*SC*) to be a function that returns true when a given number of NPs *x* is reached the cancer site *v*_*c*_ in the flow graph (*G*).

## 4 Forward Query

Forward queries (FQs) allow prediction of the biological system’s state *S* at a given time *t*_*f*_, given the initial condition in time *t*_0_ < *t*_*f*_. There are two ways to answer forward queries: using computer simulation (numerical analysis), or by analytically solving the model. Using simulation it is possible to provide as accurate results as needed for any given FQ by repeating the simulation enough times. The downside of the simulation method is the large number of repetitions required to reduce the statistical error originating in the stochastic processes of the model. In contrast, analytically solving FQ requires designing a specific algorithm for each forward query. Done right, such algorithms are less time-consuming relative to the simulation and provide accurate results. The downside is the need to design an algorithm for each FQ separately which does not scale well in the number of FQs one requires. We discuss both approaches below.

### 4.1 Simulation

Algorithm 1 is a general forward query simulation, accepting a model *M*, a current state *S*(*t*_0_), and a stop condition *SC*. It works as follows. In line 2, the variable of the simulation step is utilized with *t*_0_. In line 3, the model’s state at *t*_− 1_ is set to an empty set just to enter the main simulation loop. In lines 4-14 it executes the main simulation loop, which follows the model’s process. In line 4, we loop until the stop condition *SC* is satisfied or the model’s state does not change in the last iteration. In lines 5-11, for each node in the flow graph (*G*) we execute the interactions of the NPs between themselves and with the environment in lines 6, and 7, respectively. In lines 8-10, the NPs inside the node *v* flow to the nigher nodes *v* ∈ *N* (*v*) in size *P* (*v* → *v*_*n*_) where the NPs are picked randomly from the source node *v*_| *p*_ using the function *PickRandom* which gets a list and a size and return a random sample of the given list in the given size. In lines 12-13, we update the current model’s state and the current time to the time. In line 15, the required model’s state is returned.

#### Algorithm 1

Forward Query Simulation

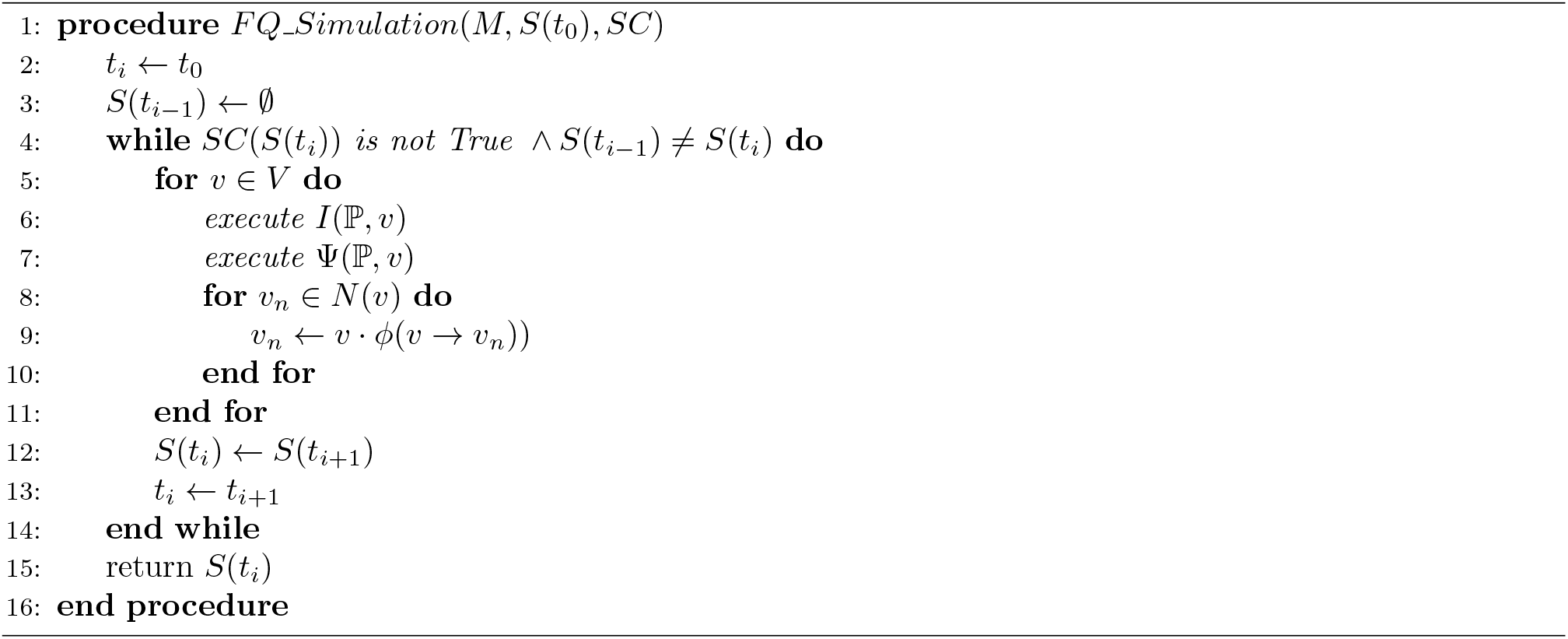

Assuming that in the worst case scenario the asymptotic complexity of *I* and Ψ are *O*(*g*(ℙ)) and *O*(*h*(ℙ)) than the simulation algorithm is performing in *O*(|*V* | · (*g*(ℙ) + *h*(ℙ + |*V* |)) per iteration. The algorithm repeats lines 4-14 a number which in the worst case is a constant number corresponding to the longest life span of a NP in the population ℙ. In each simulation iteration, three function are executed: *I*(ℙ, *v*) which assumed to be *O*(*g*(ℙ)) run time, Ψ(ℙ, *v*) which assumed to be *O*(*h*(ℙ)), and lines 8-10 which in the worst case of fully-connected graph takes *O*(|*V* |) as *N* (*v*) = *V*.

#### Lemma 1.

*If functions* Ψ *and I halts then Algorithm 1 halts*.

*Proof*. Assuming functions Ψ and *I* halts. Therefore, each algorithmic step in the main loop of the algorithm (lines 4-14 in algorithm 1) is ending after running over all the nodes in the finite flow graph.

It is sufficient to show that the condition in line 4 is not satisfies for any input at some time *t*\*. To do this, it is sufficient to show that *SC* or *S*(*t*_*i −* 1_) ≠ *S*(*t*_*i*_) is not satisfies for any input at some time *t*\*. The condition *S*(*t*_*i −* 1_) ≠ *S*(*t*_*i*_) does not satisfied where *S*(*t*_*i −* 1_) = *S*(*t*_*i*_) which happens if there is no change in the model’s state in some algorithmic step.

Recall that the model’s state contains the distribution of the population of NPs in the flow graph (*G*), where *p*_*ls*_ is the life span of a NP. There exists *t*\* = 1+max_*p*∈ℙ_ (*p*_*ls*_) such that *S*(*t*\*) = *S*(1+max_*p* ∈ ℙ_(*p*_|*ls*_)), because after the last NP in the population has exited the system, the model’s state is two zero-dimensional vectors. From this point on, the models state is always equals to a vector of zeros and in particular for *S*(*t*\*) = *S*(*t*\* + 1).□

A private case of the proposed model *M* is the *M*_*s*_ model. The *M*_*s*_ model introduce two additional condition to model *M* : 1) *I* is independent of the NPs’ type with probability *β*. 2) the NPs population’s life time in all the organs *o* ∈ *O* exponentially decays, such that each sub-population of NPs divided by their *type* has a known half life 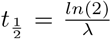 where *λ* is the decay factor. Particularly, 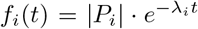 with the same 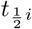 for all the organs. We define the model *M*_*s*_ as it more strictly describes the biological dynamics of *in vivo* system in rest. This is because *in vivo* experiments show that the NPs population exponentially decays in the organs and that the decay factor is depended on the type of the NPs [40, 12].

The simulation contains multiple stochastic processes. As a result, different executions of the simulator with the same arguments may provide different results. As such, one can be able to determine the model’s prediction with confidence bigger than 1 − *ϵ* for any *ϵ* ∈ [0, 1], if it runs the simulator multiple times (*n*) and computes the average output. In order to find the number of times one needs to run the simulator for given initial conditions, we compute the influence of each one of the three independent stochastic processes: 1) the interaction between the NPs and their environment Ψ in each organ node (*o* ∈ *O*) at each step in time with chance *α*. 2) the interaction between the NPs and themselves *I* at each node (*v* ∈ *V*) at each point in time. 3) the concentration of the NPs population ℙ over time, based on an exponential decay distribution. Each one of these processes would be analyzed independently in a Lemma and afterward collected together to find *n*.

Lemma 2 provides an upper boundary for the interaction between the NPs and their environment Ψ in each organ node for a single step in time.

#### Lemma 2.

*Given a M*_*s*_ *model and a FQ. It is necessary to calculated Algorithm 1’s outcome for the NPs state distribution in the organs* ∀*o* ∈ *O, n times in order to obtain a ϵ statistical error, such that n is the solution for Eq. (2) (solved for n)*.

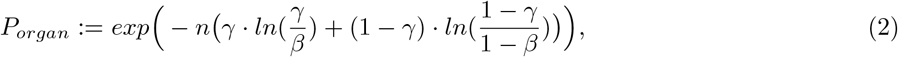

where *γ* = *β* + *E* and *β* ∈ [0, 1] is the probability that the IP *I* activates for each node (see the definition for *M*_*s*_ model).

*Proof*. For each *o*_*j*_ ∈ *O* there is a sub population of NPs at each time *t*_*i*_, marked as 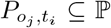. At each step in time, we perform Ψ on each NP in the organ population 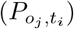. Each such interaction is an independent Bernoulli experiments with parameter *β* due to the definition of Ψ. Hence, we define *Y*_*a*_ ∼ *B*(*β*). Therefore,

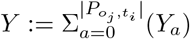

such that *Y* binomial distributed 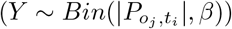.

Using Chernoff’s inequality theorem [13], each specific organ (*o*_*j*_) at each step in time *t* satisfies

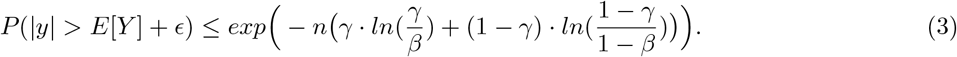

□

Therefore, an upper boundary can be taken to be

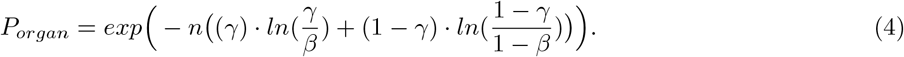

Lemma 3 provides an upper boundary for the interaction between the NPs and themselves *I* at each node (*v* ∈ *V*) for a single step in time.

#### Lemma 3.

*Given a M*_*s*_ *model and a FQ. It is necessary to calculated Algorithm 1’s outcome for the distribution in the organs and blood vessels* ∀*v* ∈ *V, n times in order to obtain a ϵ statistical error, such that n is the solution for Eq. (5) (solved for n)*.

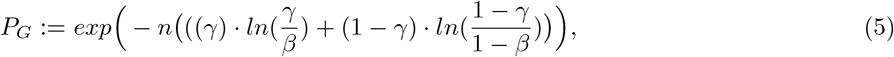

where *γ* = *β* + *ϵ*.

*Proof*. Considering *I*, for each *v* ∈ *V* there is a sub population of NPs at each time *t*_*i*_, marked as 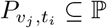. Each such interaction is an independent set of *k* Bernoulli experiments with probability *β*, following the definition of the *M*_*s*_ model. Hence, we define a *Z*_*a*_ binomial distributed variable (*Z*_*a*_ ∼ *Bin*(*k, β*)). Therefore,

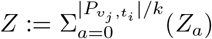

such that 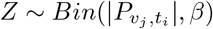.

Using the Chernoff’s inequality theorem [13], each node (*v*_*j*_) at each step in time *t* satisfies

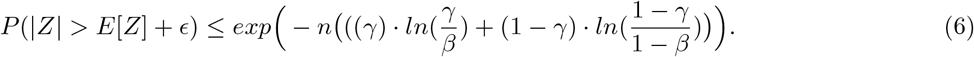

Therefore, an upper boundary can be taken to be

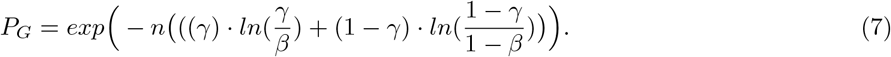

□

Lemma 4 provides an upper boundary for the concentration of the NPs population ℙ over time, based on the expected exponential concentration decay.

#### Lemma 4.

*Given a M*_*s*_ *model and a FQ. It is necessary to calculate Algorithm 1’s outcome for the population size at time t*, |ℙ (*t*)|, *n times in order to obtain a ϵ statistical error, such that n is the solution for Eq. (8) (solved for n)*.

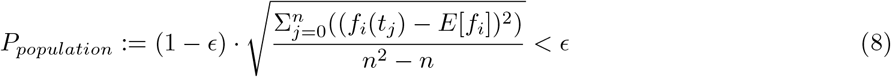

*Proof*. The concentration of the population of NP over time (|ℙ|) is dependent on the decay rate of each sub population of NPs and their initial size from the definition of *M*_*s*_ model. Therefore, mark the size of the NPs population’s distributed as Gamma-distribution with shape parameter *ρ* and inverse scale parameter *α*, Υ ∼ Γ(*ρ, α*), which satisfies

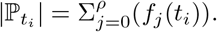

It is given by the definition of *M*_*s*_ that the sub-populations of NPs’ life span are exponential distributed with initial size |*P*_*i*_| and decay rate *λ*_*i*_ for each *i*_*th*_ sub-population of NPs. Using the Central Limit Theorem (CLT) it is possible to find an upper bound of the number of needed iterations *n*. First, it is possible to bound the required amount of samples using the wanted statistical confidence *ϵ* resulting in

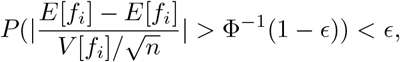

where *E*[*x*] is the mean of distribution *x* and *V* [*x*] is the standard deviation of distribution *x* and assuming the error interval is equal to the statistical error. Where, 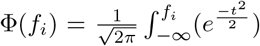 and *E*[*f*_*i*_] is the mean on the set of samples 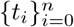. So, the stop condition is the solution to Eq. (9).

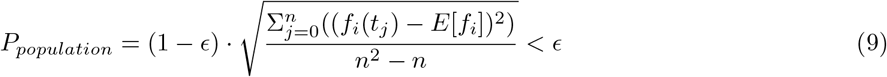

□

#### Theorem 5.

*Given model M*_*s*_ *and a forward query FQ. It is necessary to calculate Algorithm 1’s outcome for n times in order to obtain E statistical error, such that n is the solution for Eq. (10) solved for n, such that n is defined in P*_*organ*_, *P*_*G*_, *and P*_*population*_.

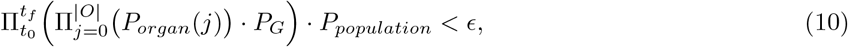

*where t*_*f*_ := *min*_*t*_(*SC* (*S*(*t*)) = 1).

*Proof*. To obtain an adequate number of times (*n*) it is sufficient to find an upper bound for each one of the processes and multiply their results to get an upper bound for the model. We use Lemmas 2-4 to obtain the upper bound of each sub-process and combine them.

From Lemma 2 it is known that for each time *t*_*i*_ and for each organ node *o*_*j*_ the upper bound satisfies Eq. (2). In addition, from Lemma 3 it is known for each time *t*_*i*_ and for each node (both organs and blood vessels) *v*_*j*_ the the upper bound satisfies Eq. (5). Because the processes are independent for each organ node and blood vessel node.

Each event is independent, so the overall upper bound is the multiplication of the individual boundaries resulting at

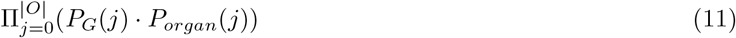

for each time *t*_*i*_. The *I*_*β*_ process is identical for each node *v*, resulting in Eq. (11) as follows:

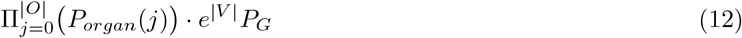

It is known that these processes are independent for each time *t*_*i*_, *t*_*j*_ such that *i* ≠ *j* and therefore, one can multiply the probabilities to obtain the boundary over time, resulting in it:

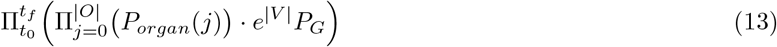

In addition, from lemma 4 it is known that the upper bound for the population size over time |ℙ (*t*)|with statistical error *ϵ* satisfies Eq. (8). Although the size of the population is directly influencing the upper boundary of the *E* and *MRIP* stochastic protocols, we relax this and assume that the processes are independent. Therefore, to obtain statistical error (*ϵ*) of model *MB* using algorithm 1, it has to be repeated *n* times, found by solving the following equation for *n*.

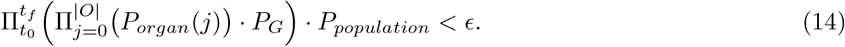

□

### 4.2 Analytical solutions for forward queries

Analytically solving FQs provides the ability to analyze the obtained results compared to FQs that are computed using a simulation which provides only an instance of the possible distribution of results. Thus, in this section, we proposed an analytical approach to answer FQs based on the proposed model by reducing each FQ into a sequence of two FQs (e.g., *FG*_1_ and *FG*_2_). Of note, the proposed method is defined for the *M*_*s*_ model rather than the *M* model (see definition in Section 4.1).

Following are two algorithms which obtains the NPs’ population distribution (ℙ) in the flow graph (*G*) at a given time *t*_*f*_, and obtains the NPs’ population’s states at a given time *t*_*f*_ marked using *FQ*_1_ and *FQ*_2_, respectively. The combination of the outputs of both algorithms, therefore, defines the model’s state at a given time *t*.

Algorithm 2, *FQ*_1_, is a general forward query obtaining the NPs’ population distribution (ℙ) in the flow graph (*G*) at a given time (*t*), accepting a model *M*_*s*_, a current state *S*(*t*_0_), and a stop time *t*_*f*_. Formally, *FQ*_1_ : (*M, S*(*t*_0_), *t*_*max*_(*t*_*f*_)) → ℕ^*n*^ where *n* = |*V* |, and *ls* stands for the life span of a given NP. This algorithms is different from Algorithm 1 as it computes just the mean distribution of the NPs population over the flow graph using a random walk approach while Algorithm 1 simulates a single stochastic outcome.

Algorithm 2 works as follows. In line 2, the function *extend organs ls* is executed which gets the model’s flow graph *G* and replace each organ node (*o* ∈ *O*) into a line graph of organ nodes with *ts* ∈ *o* nodes in the graph, each one with *ts* = 1. An example is shown in the middle view in Fig. 2. In line 3, the function *add stay nodes* is executed which for each organ node adds a new node and an edge between them with a weight *w* = *β* corresponding to the probability that Ψ ∈ *o* will make a NP stay in the organ node. An example is shown in the right view in Fig. 2. In line 4, the function *all possible paths* is executed which gets a graph, start node in the graph, and path length, and returns all possible paths in the given length, starting at the given node, as a vector. In line 5, the distribution of NPs *D* in the flow graph’s nodes (*V*) are initial to zero for each node as a mapping function between the node and the number of NPs it contains. In line 6, the paths’ weight (*PW*) is initialized with an empty vector. In line 7, the total path’s weight (*TPW*) is initialized to zero. In lines 8-12 the weight of each path in the flow graph (*G*) is calculated by summing the weight of all edges in this path (*pw*) and stored in the paths’ weight vector (*PW*) for later use. In addition, *pw* is added to the total paths’ weight (*TPW*). In line 13, the number of NPs’ which are still in the flow graph (*G*) at time *t*_*f*_ is calculated by reducing from the original population’s size at time *t*_0_ the number of NPs which left the system and marked by *SANP*. In lines 14-16, the distribution (*D*) is updated as the end of each path (marked by *path*[− 1] adds the amount of NPs that took this path to get to a specific node (*v*) in flow graph (*G*). In line 17, the algorithm returns the obtained distribution *D* and the computed paths.

**Figure 2:**
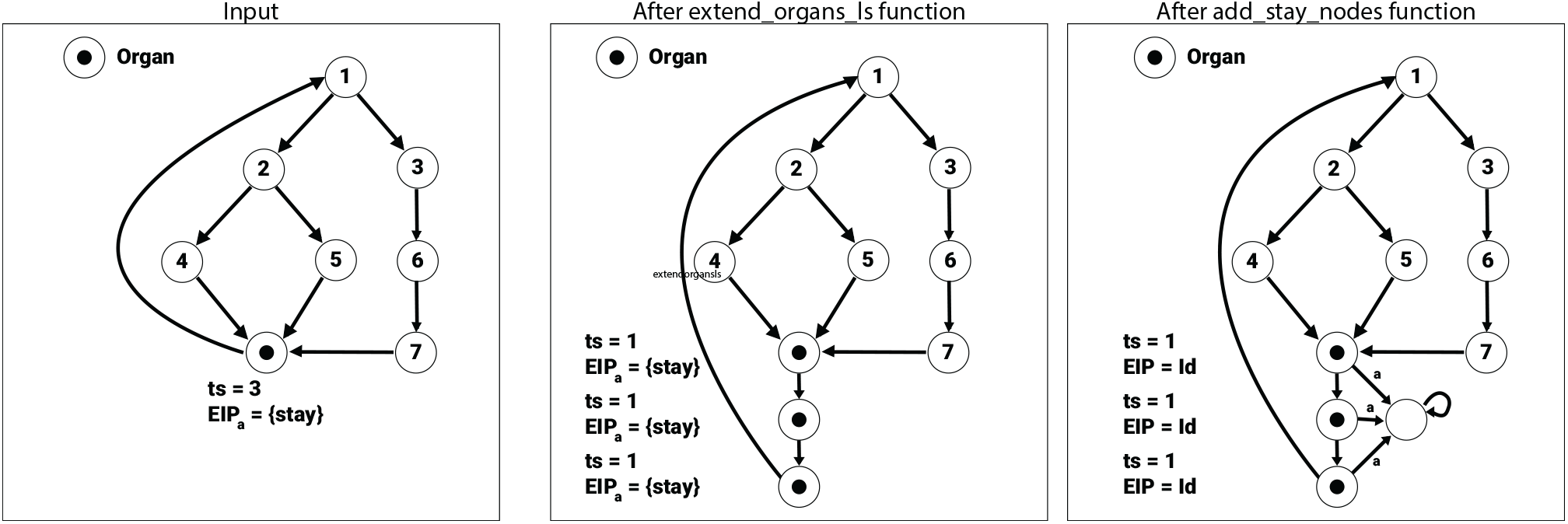
Illustration of steps *extend_organs_ls* and *add_stay_nodes* performed on a sample flow graph from algorithm 2.

#### Algorithm 2

NPs’ distribution at a given time (*FQ*_1_)

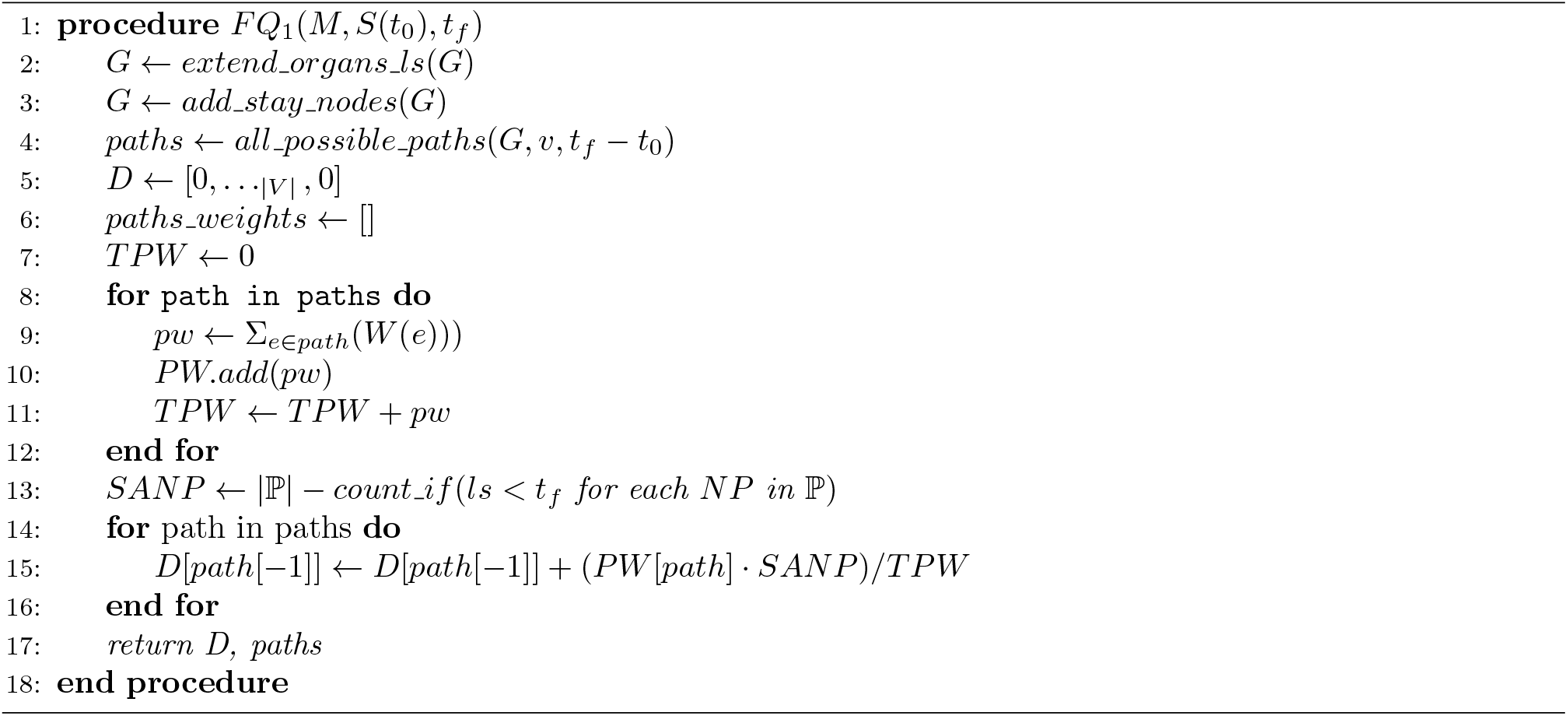

For the worst case, the asymptotic upper boundary of Algorithm *FQ*_1_’s complexity is ∀*d* > 1 : *O*(|*V* |^*d*^). where *d* is the average number of neighbors for a node *v* ∈ *V*. The performance of algorithm *FQ*_1_ is influenced by the performance of functions *extend organs ls* and *add stay nodes*. These functions can be performed by adding nodes to the adjacency list representing the flow graph (*G*) in *O*(|*O*|) time. In addition, the task of finding all possible paths with length *t*_*f*_ − *t*_0_ (in line 4) can be done using the BFS algorithm while allowing visiting nodes multiple times. The complexity of the BFS algorithm is known to be *O*(|*V* | + |*E*|). As a result, the asymptotic complexity of algorithm *FQ*_1_ is bounded by *O*(|*V* | + |*E*|) for lines 1-7. In lines 8-16, assuming there are *d* neighbors per node *v* ∈ *V* then lines 8-16 are bounded by *O*(|*V* |^*d*^) and therefore, algorithm 2 is bounded by *O*(|*V* |^*d*^) for *d* > 1. In the case of a fully connected graph the complexity is *O*(|*V* | ^|*V* |^).

Algorithm 3 *FQ*_2_ is a general forward query obtaining the NPs’ population’s state distribution at a given time, accepting a model *M*_*s*_, a current state *S*(*t*_0_), and a stop time *t*_*f*_, *FQ*_2_ : (*M, S*(*t*_0_), *t*_*max*_(*t*_*f*_)) → ℕ^|*a*|^.

It works as follows. In line 2, the state distribution is initialized to zero for all possible NP’s states. In line 3, the *FQ*_1_ algorithm is executed, providing the NP’s distribution in the flow graph (*G*) and the paths they move through. In lines 4-11, there is the main algorithmic loop that runs on each path obtained in line 3. For each organ node in the path, the Ψ function of this node is executed on the current state distribution as shown in lines 6-7. In line 8, the *I* is executed for all nodes (organs and blood vessels) in the path. In line 10, the overall NP’s states are updated according to the states obtained from this path.

#### Algorithm 3

NP’s states distribution at a given time (*FQ*_2_)

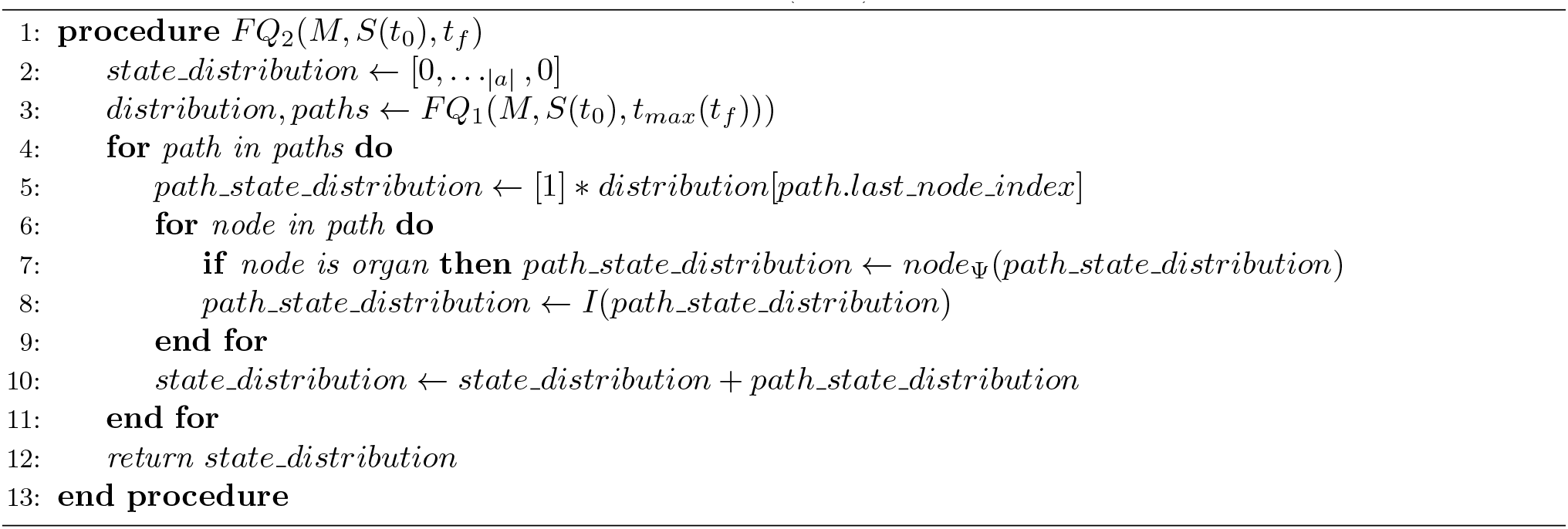

Similarly to Algorithm 2, The time complexity of algorithm 3 is bounded by *O*(|*V* |^|*V* |^) because line 3 is using algorithm *FQ*_1_ with time complexity of *O*(|*V* |^|*V* |^) and lines 3 − 12 are bounded by *O*((*t*_*f*_ − *t*_0_) · |*V* |^|*V* |^) and therefore, the overall time complexity is *O*(|*V* |^|*V* |^).

Based on Algorithms 2 and 3 (Namely, *FQ*_1_ and *FQ*_2_), it is possible to obtain the model’s state given the model (*M*_*s*_) and initial condition (*S*(*t*_0_)) at any time (*t*_*f*_). As a result, by rapidly testing if a stop condition (*SC*) is met one can obtain the answer for every forward query as shown in Lemma 6. Hence, reduces the need to explicitly develop an algorithm for each forward query separably to find a projection function from the model’s state to the needed information. In contrast, this method does not exploit the possible attributes different FQs may have and thus may not be the most optimal algorithm from a complexity point of view.

#### Lemma 6.

*From a function that executes algorithms FQ*_1_ *and FQ*_2_ *for each* 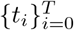, *where T* := *min*_*t*_*SC*(*S*(*t*)) *exists a reduction function to any forward query*.

*Proof*. Given a forward query *FQ*. Executing algorithms *FQ*_1_ and *FQ*_2_ on any time *t*_*i*_ provides the model’s state at this time *S*(*t*_*i*_), according to the model’s state definition. On the other hand, from Lemma 1, ∃*t*_*i*_, *i* ∈ *T* such that the condition

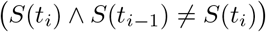

is met and therefore

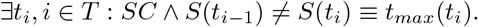

Yielding that

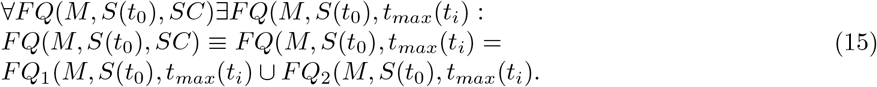

Based on Lemma 6, one can compute any FQ by computing *FQ*_1_ and *FQ*_2_ for each 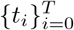 until a given stop condition *SC* defined by the user is met. Once *SC* is satisfied, one can take the obtained state *S*(*t*) and extract the sub-stable information needed. For example, if one would like to query the amount of NPs stayed in each organ after a single organ gathered *X* > 0 NPs, it could compute *FQ*_1_ and *FQ*_2_ until this condition is satisfied and then from the model’s state obtain the number of NPs in each organ node *o* ∈ *O* to answer the FQ.

## 5 Model Validation

We examine the performance of the simulation on two *in vivo* studies of NP biodistribution and their evolution over time. First, we model a mouse’s blood vessels and organs as a flow graph, using it as the graph *G* of a *M*_*s*_ model. This model is defined as the *baseline* instance of the simulation. The *baseline* instance will be modified for each *in vivo* study, by replacing the NP’s population half-life, injection node, and Ψ in the organs. The *baseline* instance is implementing the *M*_*s*_ model (see Section 4.1) since the NPs’ concentration decay exponentially and the interaction protocol between the NPs (*I*) assumed to be the identity function.

### 5.1 Model mouse flow graph construction

The flow graph used in the simulator is based on research by Aslanidou et al. [7] which describes the cardiovascular system’s topology of a mouse with 84 blood vessels and the heart. Specifically, the radius, average blood pressure, and length of each blood vessel.

We constructed a flow graph so that each blood vessel is represented by *i* ∈ ℕ blood vessel nodes, where *i* is the length of the blood vessels in millimeters (mm) (rounded to the closest natural value) forming a line-graph where the weights of the edges between the nodes are the radius of the blood vessel in mm. For example, the *ascending aorta* blood vessel is reported to be with a length of 2.6 mm and with an average radius of 0.705 mm so there will be 3 blood vessel nodes with a weight of 14100(= 0.705 ∗ 2 ∗ 1000) (representing the maximal number of NPs that can pass in a single clock tick at once) between each two neighbor nodes.

In addition, six more organs (spleen, left and right kidneys, liver, and left and right lungs) have been introduced to the flow graph according to the weight of the organs, and blood volume based on a study by Šebestik et al. [54]. The organs connected to the cardiovascular system (described in [7]) using the schematic connection described in [14]. This results in a flow graph with seven organs (spleen, left kidney, right kidney, liver, left lung, right lung, and heart) in total. Two of them (lungs and kidney) have both left and right instances which results in five distinctive organs. The model contains seven organ nodes, 934 blood vessel nodes, and 995 edges.

The tissue microenvironment interaction protocol Ψ of all introduced organs makes the NPs stay in the organ with a given probability *c*. In addition, the time span (*ts*) of each organ is set according to the blood vessels capacity of the organ [54]. A mouse’s blood volume is approximately 21.8 milliliter (ml) on average [54]. Assume there are only seven organs according to the mouse’s anatomy and blood vessels (as described by [7]), the sum of the edges between all the blood vessels is exactly the volume of the blood in the mouse’s body without the average blood volume in the organs. Therefore, the average time span, *ts*_1_, for a single blood vessel node is defined by

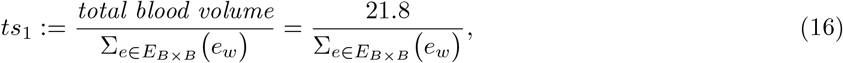

where *E*_*B* ×*B*_ ⊂ *E* is the set of edges such that the source and target nodes of an edge are blood vessel nodes. *e*_*w*_ is the weight of edge *e*. The average weight, blood volume and time span (*ts*) for each organ are summarized in Table 1. The first row indicates the average *in vivo* measurement of each organ’s weight in grams and the second row indicates the average blood volume in millimeters. The third row indicates the average number of model’s clock tics a NP spend in an organ, computed using the formula 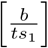 in order to keep the total blood volume from Eq. (16) the same.

**Table 1:** The weight, blood volume, and the time span for each type of organ in the mouse used in the simulation. * The value was calculated using linear regression (see Eq. (17)).

|  | Heart | Kidney | Lung | Liver | Spleen |
| --- | --- | --- | --- | --- | --- |
| Average organ weight (g) [54] | 1.0 | 2.5 | 2.8 | 13.1 | 0.9 |
| Average organ blood volume (ml) [54] | 0.24* | 0.67 | 0.78 | 3.68 | 0.19 |
| Organ's time span (model's clock ticks) | 7 | 17 | 19 | 90 | 6 |

Since the heart’s blood volume is not provided by [54], we assume the relationship between the other organs’ weight and blood volume is kept and linear. Thus, we computed a linear regression using the data of the kidney, lung, liver, and spleen obtaining the following relationship:

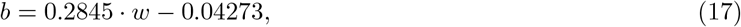

such that *b* is an organ’s blood volume and *w* is an organ’s weight. The linear model (Eq. (17)) obtained with coefficient of determination of *R*^2^ = 0.9998, indicating that the relationship between organ’s weight and blood volume is indeed linear. From [54], we know that the mouse’s average wright is 1.0 grams. Hence, by setting *w* = 1.0 in Eq. (17) one obtains the heart’s blood volume to be 0.24.

A 2D representation of the flow graph is shown in Fig. 3 where the yellow dots are the blood vessel nodes *B*, the black dots are organ nodes *O*, and the lines are the directed connections between them (*E*) such that the thickness is linear relative to the edge’s weight.

**Figure 3:**
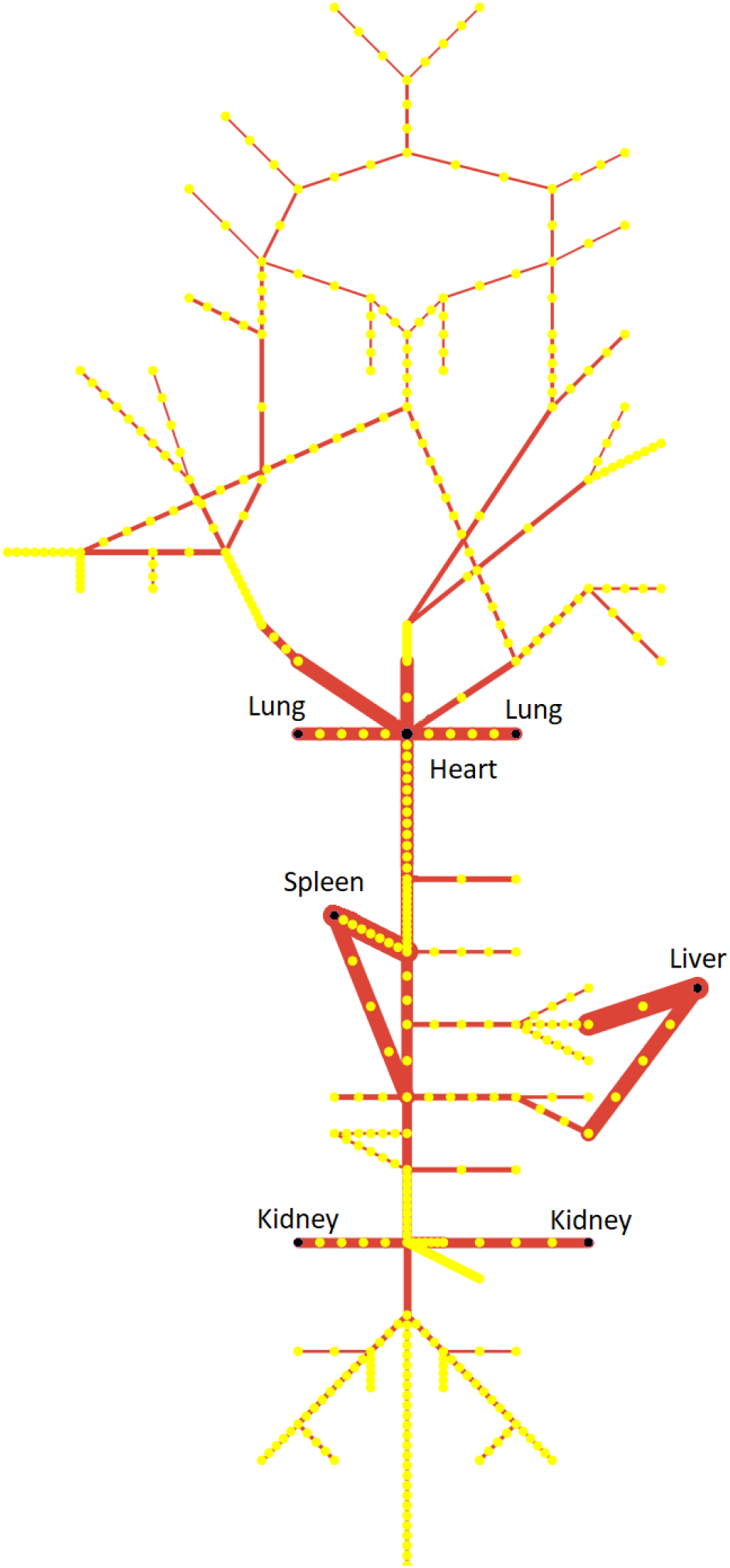
2D illustration of mouse’s cardiovascular system model as used in the simulation based on [7, 14]. The yellow dots are the start and end of blood vessels (*B* ∈ *G*), the purple dots are organs (*O* ∈ *G*), and the red arrows are the directed connections between them (*E* ∈ *G*).

### 5.2 Experiment setup

In order to evaluate the model’s ability to accurately predict the PKPD dynamics of an NP-based drug in general and the targeted drug delivery capabilities in particular we conducted a series of experiments. In each experiment, the simulator is configured according to the data of the *in vivo* experiment, predicted the experiment’s (i.e., the *in silico*) results, and compared with the corresponding *in vivo* results. High correlation between the *in silico* and *in vivo* results strongly indicates the ability of the model to replace *in vivo* experiments up to some limit.

The model’s parameters used in the following experiments were calculated based on *in vivo* data. Later, for each experiment, several parameters will be modified to better represent the specific data related to each experiment. For example, the half-life of the NPs’ population is a direct attribute of the type of the NP and therefore be modified according to the NP properties used in the experiment.

A population of 50000 NPs of the same type ℙ was used for the simulation each time. The population size of 50000 is picked as in the flow, the amount of NPs is directly influencing the distribution and therefore a small population can introduce additional errors due to the calculation with integers. A bigger population may result in a better precision while calculating Eq. (1). On the other hand, a larger population increases the computation time. An NP population of size 50000 was picked empirically to balance these two constraints.

The model uses abstract discrete time steps. In order to calibrate the simulation’s abstract time step into a real one, we find the duration of a model’s clock tick, based on Table 2, as follows. The four blood vessels (ascending aorta, thoracic aorta, right renal, and left renal) are presented in the flow graph of the simulator (Fig. 3) were taken and are summarized in Table 2. The average blood flow velocity 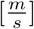 and respectively blood vessels length [*m*] were taken from Aslanidou et al. [7]. The average flow velocity divided by the length of the blood vessels is approximately 8.03 · 10^−5^ seconds which is set to be the model’s clock tick duration.

**Table 2:** Calibration of the model’s discrete clock tick duration based on *in vivo* data of the average time it would take an NP to cross a single blood vessel node. The model clock tick duration obtained from each sampled blood vessel separately results in a slightly different value due to errors in the physical and biological properties’ measurements and in the numerical calculation.

| Blood vessel name | Ascending aorta | Thoracic aorta | Right renal | Left renal |
| --- | --- | --- | --- | --- |
| Average blood flow velocity $[\frac{m}{s}]$ | $1.7 \cdot 10^{-1}$ | $9.0 \cdot 10^{-2}$ | $1.1 \cdot 10^{-1}$ | $1 \cdot 10^{-1}$ |
| Blood vessel Length $[m]$ | $2.6 \cdot 10^{-3}$ | $1.26 \cdot 10^{-2}$ | $2.9 \cdot 10^{-3}$ | $1.5 \cdot 10^{-3}$ |
| Passing time $[s]$ | $4.42 \cdot 10^{-4}$ | $1.13 \cdot 10^{-3}$ | $3.19 \cdot 10^{-4}$ | $0.015 \cdot 10^{-4}$ |
| Blood vessels nodes in model | 4 | 14 | 4 | 3 |
| Model clock tick $[s]$ | $1.11 \cdot 10^{-4}$ | $8.10 \cdot 10^{-5}$ | $7.98 \cdot 10^{-5}$ | $5.00 \cdot 10^{-5}$ |

The simulation (Algorithm 1) was executed rapidly according to Theorem 5 for each type of NP with allowed statistical error of *ϵ* = 0.05 for each type, resulting in around 10000 repetitions for each NP. The number of repetitions is slightly different between NP’s types due to each population type’s half-life duration.

The comparison between the *in silico* results obtained in each experiment and the *in vivo* values (Sections 5.3 and 5.4) is presented using a scatter plot where the x-axis is the *in vivo* value and the y-axis is the *in silico* result. The values in the *in vivo* experiments are normalized, resulting in the distribution of NPs in the organs. Therefore, the graph is bounded by [0, 1] × [0, 1]. Due to the normalization of the values, the cases where the simulator predicts exactly the *in vivo* values or returns random values basically define the best and worst cases, respectively.

Where the simulator predicts exactly the *in vivo* values, the dots will arrange on the line *in silico* = 1 · *in vivo* + 0, each *in vivo* value is equal to the corresponding *in silico* result as shown in Fig. 4 (dotted line). Alternatively, where the simulation returns values that are independently of the *in vivo* values (i.e. uniformly distributed), the dots arrange on the line *in silico* = 0 · *in vivo* + 1*/* |*O* |. Because the chance an NP in the population will be allocated to each organ is the same. Therefore, after normalization, each organ will have 1*/* |*O* | NPs as shown in Fig. 4 (solid line).

**Figure 4:**
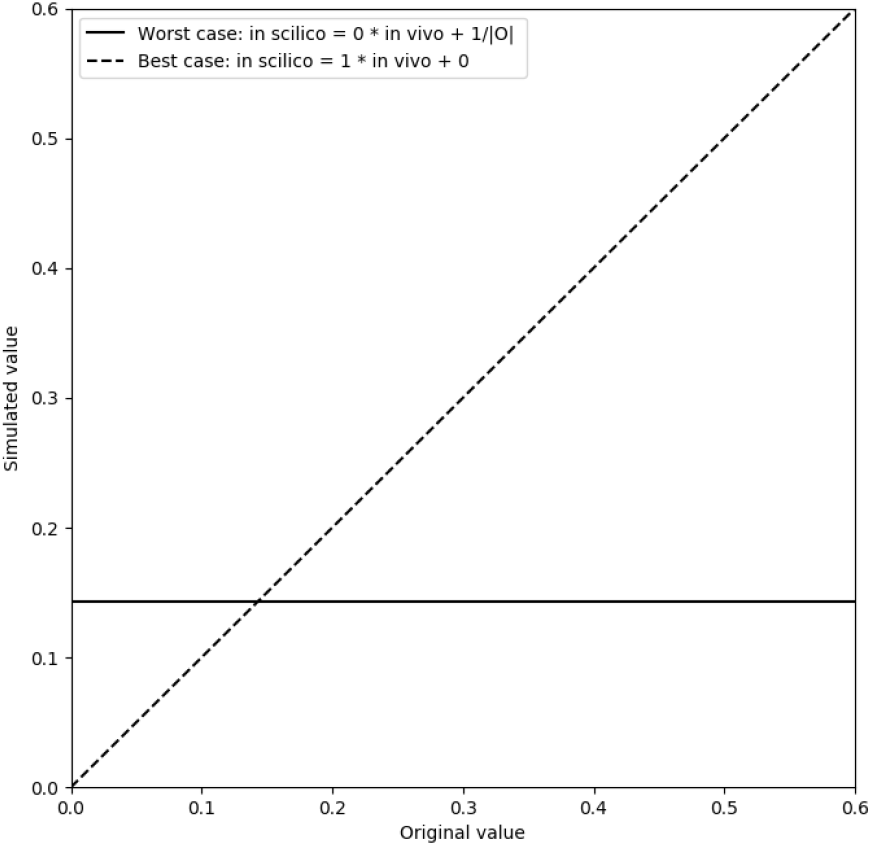
The solid and dashed lines indicate the optimal and worst predict assessments of the model’s *in silico* results in comparison to the *in vivo* values.

As a result, by comparing the line obtained using linear regression of the dots (*in vivo, in silico*) to the ideal line one can compare the performance of the simulator. Specifically, as the *x*-intercept of the linear regression line is smaller and the slope is closer to 1, the better the model predicts the *in vivo* values.

### 5.3 Biodistribution of shape and red blood cell (RBC) membrane depended on nanoparticles in mice

Ben Akiva et al. [12] investigated the performance of six types of NPs as a targeted drug delivery platform *in vivo* on mouse. The six types of NPs are described in Table 3. All NPs are made of lactic-co-glycolic acid (PLGA) and their size range between 200 and 240 nanometers (nm) in diameter on average [12].

**Table 3:**
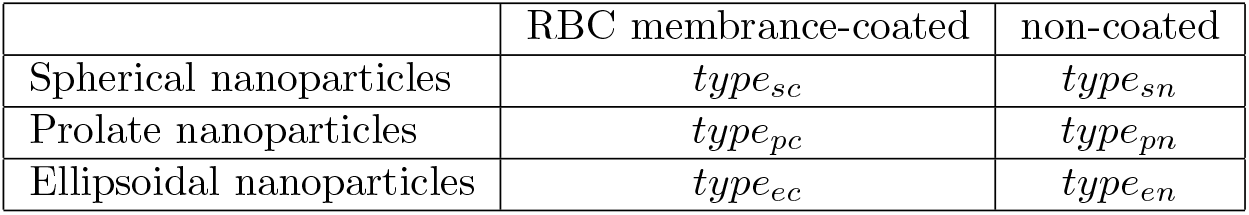
The six different types of NPs used in [12].

The authors investigate how coated and non-coated red blood cell (RBC) NP biodistribution is affected by their shape (spherical, prolate, and ellipsoidal) in mice. They injected the NPs using retroorbital injections and sampled the NP’s biodistribution after 0.25, 0.5, 0.75, 2, 4, and 24 hours. They found that all six types of NP populations decay exponentially [12]. The respective half-life of each type of NP population has been calculated and presented in [12]. 24 hours after the NPs injection, the mice were euthanized and the spleen, kidney, liver, lungs, and heart were dissected and imaged. Then, the normalized distribution of the NPs in these organs was calculated. The values from [12] were taken directly from the plot using Adobe Illustrator software by calculating the height of the columns in pixels. The sum of all heights of rectangles in the same color (representing the biodistribution of one type of NP) was summed up to the y-axis’s height and obtained with an average error for an NP’s type is 0.4%. The simulation (Algorithm 1) executed for all six types of NPS shown in Table 3.

### 5.4 Biodistribution of cholorotoxin-conjugated iron oxide nanoparticles in mice

Lee et al. [40] investigated the performance of two types of NP (NP-CTX-chitosan and NP-chitosan) which are made of iron oxide coated with chitosan and polyethylene glycol with a mean diameter of 7 nm for targeted drug delivery. The authors investigate how to avoid synthesizing radioactive NPs and evaluate them in mice in PK models by assessing serum half-life and tissue distribution of NPs in mice [40]. The authors injected the NPs using the tail vein and sampled the NP’s biodistribution after 1, 3, 6, 10, 24, and 48 hours. Similar to [12], Lee et al. found that the half-life of both NPs types decay exponentially [40].

In addition, after 6, 24, and 48 (3 mice for each case) the mice were euthanized and 12 organs were dissected and imaged. From these images and a baseline image of the fluorescence, the authors calculated the average fluorescence intensity per organ. Assuming the amount of fluorescent material each NP caring is the same, the results are linearly correlated to the amount of NPs in each organ. Lee et al.’s results are presented in [40].

This simulation differs from the simulation presented in Section 5.3 by the following three parameters. First, the injection node is the tail vein rather than the retroorbital vein. Second, the half-life of NP-CTX-chitosan and NP-chitosan is assumed to be 8 and 7 hours, respectively [40]. Third, We assign the probability a NP stays in an organ *c* to be 0.02 after simulating the results for *c* ∈ [0.01, 0.015, 0.02, 0.025] and comparing the slope of the linear regression model upon the data, getting best results for *c* = 0.02. We tested larger *c* as the NPs’ size is two magnitudes smaller than the ones used by Ben-Akiva et al. [12] (7nm compared to 200nm-240nm) and therefore have a larger chance to stay in an organ.

## 6 Results

### 6.1 A comparison between in vivo and in silico results of the biodistribution of RBC nanoparticles with different shapes

Comparison between the simulation (*in silico*) results and the *in vivo* values from ([12] Fig. 4C) is shown in Fig. 5, where each dot is the average values of *n* = 70 and the error bars are one standard deviation error. The NPs’ types and the organs are represented in the graph by dots in different colors and shapes, respectively. The solid (gray) line shows the ideal linear regression for comparison. A linear regression has been performed on the tuples (*in vivo, in silico*),

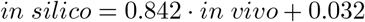

with coefficient of determination *R*^2^ = 0.933 and shown as a dashed (black) line. The area with 95% confidence level of the model is marked in gray [17]. From an error analysis of the model’s parameters, we obtain that the linear regression’s slope, intercept, and *R*^2^ are 0.842 ± 0.01, 0.032 ± 0.002, and 0.933 ± 0.004 (presented as mean ± standard deviation), respectively.

**Figure 5:**
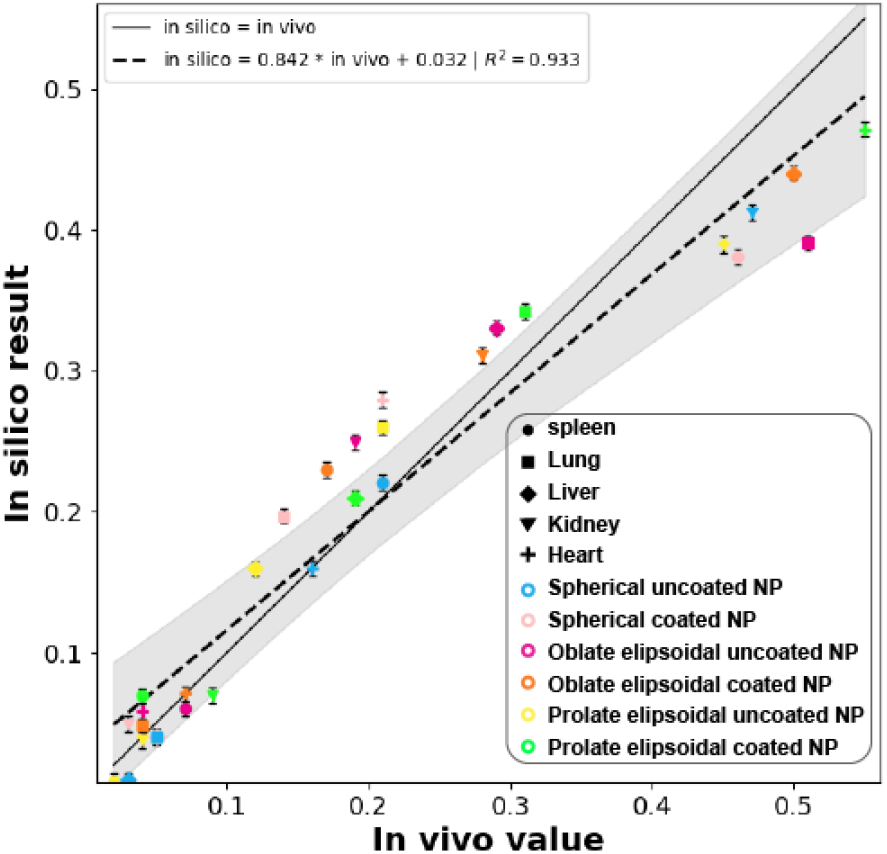
Comparison between the *in vivo* and *in silico* results for the relative amount of each type of NPs located in each organ after 24 hours. The *in vivo* research was conducted by Ben-Akiva et al. [12] and compared to our *in silico* calculations using the simulator. The line is a result of the linear regression on the presented values with coefficient of determination *R*^2^ = 0.933. For each dot, the shape indicates the organs and the color the type of the NPs.

A breakdown of the graph (Fig. 5) into each of the six types of NP’s is presented in Table 4 as a linear regression of dots. In a similar manner, Table 5 shows the breakdown of Fig. 5 into the five organs. These results show that the simulator better predicts the biodistribution of NPs inside the body compared to the predictions of several NPs in a single organ as the average fitting for the NPs is 0.844 while for the organs is 0.653. A possible explanation for this outcome is the fact, that the biodistribution of NPs in the body across the organs is normalized which eases the task. Specifically, for the lungs, the simulator obtains 0.357 fitting to the *in vivo* values which decreases the average score significantly. This is consistent with the results obtained by Ben-Akiva et al. that obtain *p* > 0.1 on the lungs-related data for their experiment on three mice [12].

**Table 4:** Fitting between the simulator’s (*in silico*) results and Ben-Akiva et al. [12] *in vivo* values for each NP’s type, shown as (mean ± standard deviation).

| NPs type | Linear regression slope | Coefficient of determination $R^2$ |
| --- | --- | --- |
| Spherical coated | $0.845 \pm 0.017$ | $0.968 \pm 0.006$ |
| Spherical uncoated | $0.713 \pm 0.017$ | $0.886 \pm 0.013$ |
| Oblate ellipsoidal coated | $0.935 \pm 0.018$ | $0.945 \pm 0.007$ |
| Oblate ellipsoidal uncoated | $0.877 \pm 0.020$ | $0.945 \pm 0.008$ |
| Prolate ellipsoidal coated | $0.804 \pm 0.019$ | $0.950 \pm 0.008$ |
| Prolate ellipsoidal uncoated | $0.892 \pm 0.021$ | $0.907 \pm 0.012$ |
| **Average** | $0.844 \pm 0.0187$ | $0.9335 \pm 0.009$ |

**Table 5:** Fitting between the simulator’s (*in silico*) results and Ben-Akiva et al. [12] *in vivo* values for each organ, shown as (mean ± standard deviation).

| Organ | Linear regression slope | Coefficient of determination $R^2$ |
| --- | --- | --- |
| Heart | $0.813 \pm 0.101$ | $0.652 \pm 0.098$ |
| Kidney | $0.617 \pm 0.075$ | $0.449 \pm 0.075$ |
| Liver | $0.714 \pm 0.071$ | $0.581 \pm 0.064$ |
| Lung | $0.357 \pm 0.147$ | $0.110 \pm 0.102$ |
| Spleen | $0.766 \pm 0.048$ | $0.952 \pm 0.023$ |
| **Average** | $0.653 \pm 0.0948$ | $0.6586 \pm 0.0724$ |

From these results and Table 6, it is possible to see that as the population of NPs’ half-life is larger, the *in silico* results better fits the *in vivo* values. One possible explanation is that a longer half-life means NPs stay longer in the system and it is easier to predict their biodistribution. Indeed, the worst predictions have been obtained for the NP’s type with the shortest half-life.

**Table 6:** The connection between the population of NPs’ half-life to the simulation fitting to the *in vivo* experiment.

|  |  |  |  |  |  |  |
| --- | --- | --- | --- | --- | --- | --- |
| Half-life (minutes) [12] | 1476 | 3888 | 3960 | 4560 | 4920 | 10296 |
| Fitting (mean) | 0.713 | 0.845 | 0.877 | 0.892 | 0.804 | 0.935 |

### 6.2 Biodistribution of cholorotxin-conjugated iron oxide NP in mice

Comparison between the simulation’s (*in silico*) results and the *in vivo* values from [40] is shown in Fig. 6, where each dot is the average values of *n* = 70 repetitions of the simulator and the error bars are one standard deviation error. The NPs’ types and the different organs are represented in the graph by dots in different colors and shapes, respectively. The gray (solid) line shows the ideal linear regression for comparison. A linear regression has been performed on the tuples (*in vivo, in silico*),

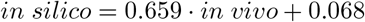

with coefficient of determination *R*^2^ = 0.946 and shown as a dashed (black) line. The area with 95% confidence level of the model is marked in gray [17]. From an error analysis of the model’s parameters, we obtain that the linear regression’s slope, intercept, and *R*^2^ are 0.659 ± 0.012, 0.068 ± 0.002, and 0.944 ± 0.008 (presented as mean ± standard deviation), respectively.

**Figure 6:**
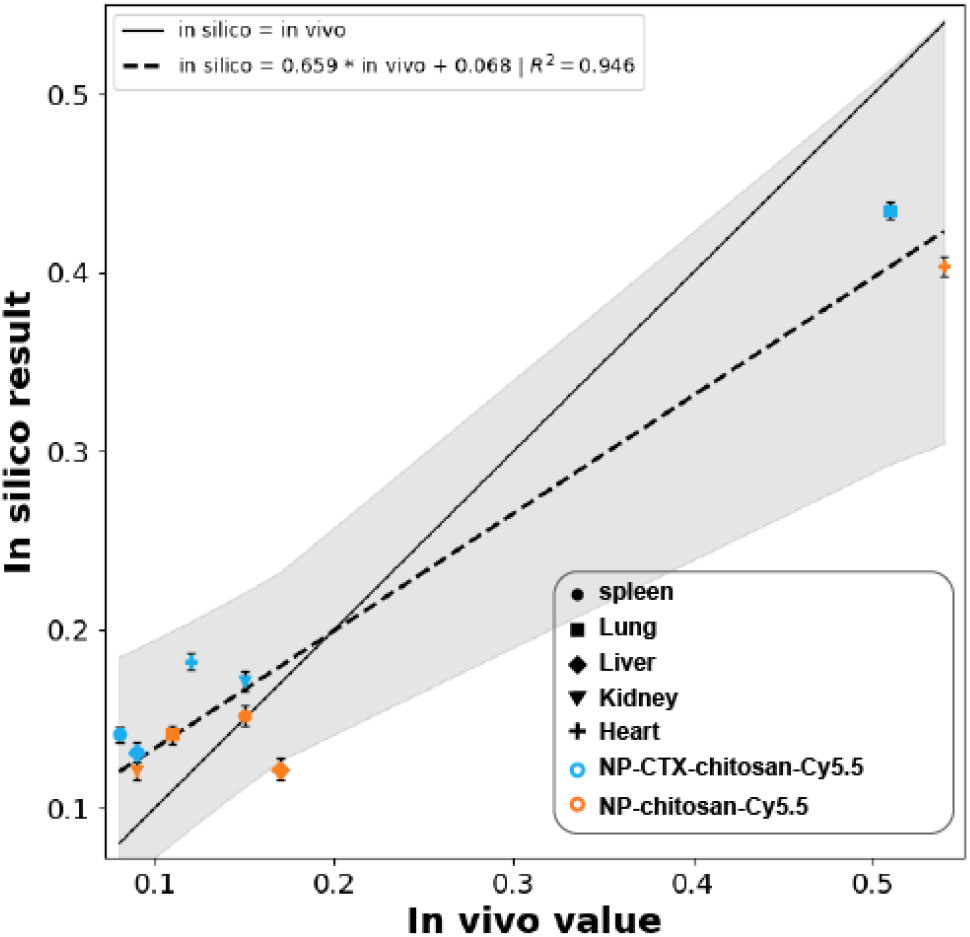
Comparison between the *in vivo* amount of NPs stayed in each organ type after 24 hours as presented by Lee et al. [40] (on the x-axis) and the *in silico* amount calculated using the simulator (on the y-axis). The line is a result of the linear regression on the presented values with coefficient of determination *R*^2^ = 0.946. For each dot, the shape indicates the organ and the color the type of the NP.

It is possible to see that except for the liver, the other four organs have a similar amount of NP. Compared to the NPs used in [12], the NP used for this experiment has a much higher half-life time (7, 8 hours) and better chance to accumulate in organs [40]. As a result, we obtain a more equally distributed biodistribution of the NPs in the organs. Which, as a result, produces 0.659 fitting.

A breakdown of Fig. 6 into the two NP’s types shows that the linear regressions based on the NP-CTX-chitosan and NP-chitosan biodistributions are *in silico* = 0.715 · *in vivo* + 0.055 and *in silico* = 0.597 · *in vivo* + 0.077 with coefficient of determination *R*^2^ = 0.938 and *R*^2^ = 0.974, respectively.

## 7 Discussion

This study presents a novel PKPD mathematical model for NP-based drugs. The model takes into consideration the biodistribution and the NPs’ population’s state’s distribution, as well as providing a platform for single-type NPs’ swarm. Based on the proposed model (see Section 3), we provide a method to numerically and analytically perform prediction (forward) queries.

We evaluate the proposed model using two *in vivo* experiments which conducted by Ben-Akiva et al. [12] and Lee et al. [40], obtaining 0.842 ± 0.01 with a coefficient of determination *R*^2^ = 0.933 ± 0.004 and 0.659 ± 0.012 with *R*^2^ = 0.946 ± 0.008, as shown in Figs. 5 and 6, respectably. An evaluation of the simulator’s results compared to the *in vivo* values from [12] for each of the six NPs, as presented in Table. 3, indicates that the model has a mean accuracy of 0.844 (±0.0187). Similarly, for the same experiment, dividing the results according to the five organs (heart, kidney, liver, lung, and spleen) obtained to be 0.653 ± 0.0948, as shown in Table. 5. Thus, the model fairly predicts the biodistribution of the NPs in the mouse’s body. The model accuracy is insufficient for analyzing an individual organ with several NPs. One possible explanation for this result could be that different types of NPs have different interaction protocols with the tissue’s microenvironment of each organ. However, due to the lack of relevant biological data, during the simulation, we assumed that the interaction protocol is identical for all types of NPs and organs.

Moreover, the flow graph is based on the research of Aslanidou, Trachet, Reymond, Fraga-Silva, Segersm, and Stergiopulos [7] and Bi, Deng, Murry, and An [14]. Aslanidou, Trachet, Reymond, Fraga-Silva, Segersm, and Stergiopulos [7] report the blood vessels length and average radius for different varieties of mice and Bi, Deng, Murry, and An [14] the connection to of the cardiovascular system to other organs. As a result, the integration between them introduces error in the blood vessels leading to the spleen, liver, and kidneys from the heart. A more detailed vascular system graph should significantly improve the simulator results. Furthermore, introducing of additional organs to the simulation will generate a analysis of the PKPD dynamics.

Finally, we assume a simple flow dynamics of the NPs’ population in the flow graph (Eq. (1)) which provides an approximation to the real NPs’ flow in the blood. On one hand, this approach introduces approximation error to the model but on the other hand keeps the calculations manageable and stable, picking one side of the well-known trade-off between fast and stable numerical calculation and accuracy. On the other side of the trade-off, one can solve the Navier-Stokes equation for each blood vessel independently to achieve a high level of accuracy [44, 6, 27]. Furthermore, this approximation works well on a large scale and long processes such as in this research. In future work, we plan to use a detailed flow dynamics model to improve the accuracy of the model.

Based on the proposed model, one can define a backward query. A backward query can be used to obtain what action should be taken at the beginning of the treatment (e.g. *t*_0_) to obtain some desired state at a given time (*t*_*f*_) in the future. An example of a backward query is: given a desired biodistribution of the NPs’ population at the end of the treatment, to which blood vessels and what subgroups of the NPs’ population should be injected into the body at the beginning of the treatment. A possible definition of the backward query is a function *BQ* such that *BQ* : (*M, S*(*t*_*f*_)) → 𝕊 which satisfies ∀ℙ ∈ **P**_*s*_ : *FQ*(*M, S*(*t*_0_), *t*_*max*_(*t*_*f*_)) = *S*(*t*_*f*_), where *M* is the model, 𝕊 is the space of all possible tuples (*S*(*t*_0_), **P**_*s*_) where *S*(*t*_0_) is the state of the model at time *t*_0_ (e.g. after injection), *S*(*t*_*f*_) is the state of the model at time *t*_*f*_ where *f* ∈ *N*, **P**_*s*_ ⊂ ℙ where ℙ is the space of all possible NP populations **P**, and *t*_*max*_(*t*_*f*_) is a function which returns 0 if *t* < *t*_*f*_ and 1 otherwise. The analysis of the backward query on the proposed model and developing an efficient algorithm for calculating it is a possible extenuation of this research.

## Notes

### Competing Interest Statement

The authors have declared no competing interest.

